# Formation and Spontaneous Long-Term Repatterning of Headless Planarian Flatworms

**DOI:** 10.1101/2021.01.15.426822

**Authors:** Johanna Bischof, Jennifer V. LaPalme, Kelsie A. Miller, Junji Morokuma, Katherine B. Williams, Chris Fields, Michael Levin

## Abstract

Regeneration requires the production of large numbers of new cells, and thus cell division regulators, particularly ERK signaling, are critical in regulating this process. In the highly regenerative planarian flatworm, questions remain as to whether ERK signaling controls overall regeneration or plays a head-specific role. Here we show that ERK inhibition in the 3 days following amputation delays regeneration, but that all tissues except the head can overcome this inhibition, resulting in headless regenerates. This prevention of head regeneration happens to a different degree along the anterior-posterior axis, with very anterior wounds regenerating heads even under ERK inhibition. Remarkably, 4 to 18 weeks after injury, the headless animals induced by ERK inhibition remodel to regain single-headed morphology, in the absence of further injury, in a process driven by Wnt/β-catenin signaling. Interestingly, headless animals are likely to exhibit unstable axial polarity, and cutting or fissioning prior to remodeling can result in body-wide reversal of anterior-posterior polarity. Our data reveal new aspects of how ERK signaling regulates regeneration in planaria and show anatomical remodeling on very long timescales.

## Introduction

Following injury or disease, some animals are capable of replacing lost structures^1^. Such regenerative processes require the production of new cells, in most systems achieved through the activation of stem cells^2,3^. ERK/MAP Kinase signaling is one of the crucial regulators of cell division^4^ and has been shown to play an important role in many model systems of regeneration^5,6^. In the highly regenerative planarian flatworm, ERK phosphorylation is one of the first responses to injury, detectable 15 min after cutting^7^. ERK signaling has been argued to be important in planaria for overall regeneration^7^, to play a role in allowing differentiation^6^, to activate stem cell subpopulations^8^ and to play a head-specific role^9^. Several studies have shown that ERK inhibition immediately following amputation leads to the formation of headless animals^6,7,9^.

The variety of regenerative activities that ERK signaling has been reported to mediate suggests that ERK signaling exists in a complex milieu of different signaling factors that interact in ways that are only beginning to be understood^10^. For example, Follistatin, a secreted inhibitor of activin signaling, was recently shown to be conditionally required for head regeneration in planaria, dependent on the underlying Wnt signaling environment at the injury site^11,12^, illustrating that the complex factors that regulate regeneration are interacting in a dynamic and spatial distinct manner that remains to be fully described.

Planarian regeneration is a complex and highly regulated process, that is also very dynamic, with planarians completing regeneration within 7-10 days, regardless of whether they are forming a wild-type anatomy or heteromorphoses, such as 2-headed or headless morphologies^13^. Even in the absence of major wounding events, planaria continuously adjust their size to changes in nutritional availability by shrinking and growing to maintain correct proportions^14,15^. Underpinning this size adaptation is the constant cell turnover which occurs around every 30 days, the longest regenerative timeframe that has been described in planaria^16,17^. Most regeneration studies in planaria stop characterizing regeneration effects around 14 days after injury and little work has explored repatterning and changes to morphology on longer timescales. Planarian regenerative outcomes, such as the headless worms induced by ERK inhibition, have previously been assumed to be permanent^7^.

Here we show that ERK inhibition for the first 3 days after amputation specifically inhibits head regeneration, while other tissues, such as the tail, can fully regenerate after ERK inhibition. This effect of ERK inhibition on regenerative outcomes is dependent on the underlying signaling environment, with head regeneration still occurring following ERK inhibition in the very anterior regions of the animal. Remarkably, we found that without any further external manipulation, the headless animals generated through ERK inhibition do not permanently remain headless but spontaneously begin to repattern to a wild-type anatomy anywhere between 4 and 18 weeks after their original formation. This repatterning occurs after the regeneration-driven remodeling is complete and is regulated by Wnt/β-catenin signaling. Finally, we show that headless animals appear to lack stable axial polarity, as their anterior-posterior axis can become reversed following bisection or fissioning, with a head forming from a posterior blastema. Thus, our findings reveal that *D. japonica* can undergo spontaneous repatterning to overcome the headless morphology and remodel tissue long after regeneration is completed, indicative of remodeling processes controlling morphology on longer timescales than previously known.

## Result and Discussion

### Short term ERK inhibition selectively blocks regeneration of anterior tissue

ERK signaling has been shown to play a role in controlling regeneration in both *Schmidtea mediteranea* and *Dugesia japonica* planarians^6,7,9^. Therefore, with the goal of exploring the effect of a short-term inhibition of regeneration on overall regenerative outcomes in *Dugesia japonica*, we treated pre-tail fragments (with both anterior and posterior wounds requiring regeneration), for 3 days with the highly specific and efficient ERK inhibitor U0126^7^ (Figure 1A). Following this 3-day treatment, we found that 98% of fragments regenerated as headless animals within 7 days (Figure 1B), lacking both external head structures and a brain (Figure 1B). The regeneration was determined to be complete by closure of the wound, pigmentation of any blastema tissue and formation of a stable morphology. This recapitulates previous reports that various ERK inhibition methods all result in headless animals^6,7,9^.

**Figure 1:**
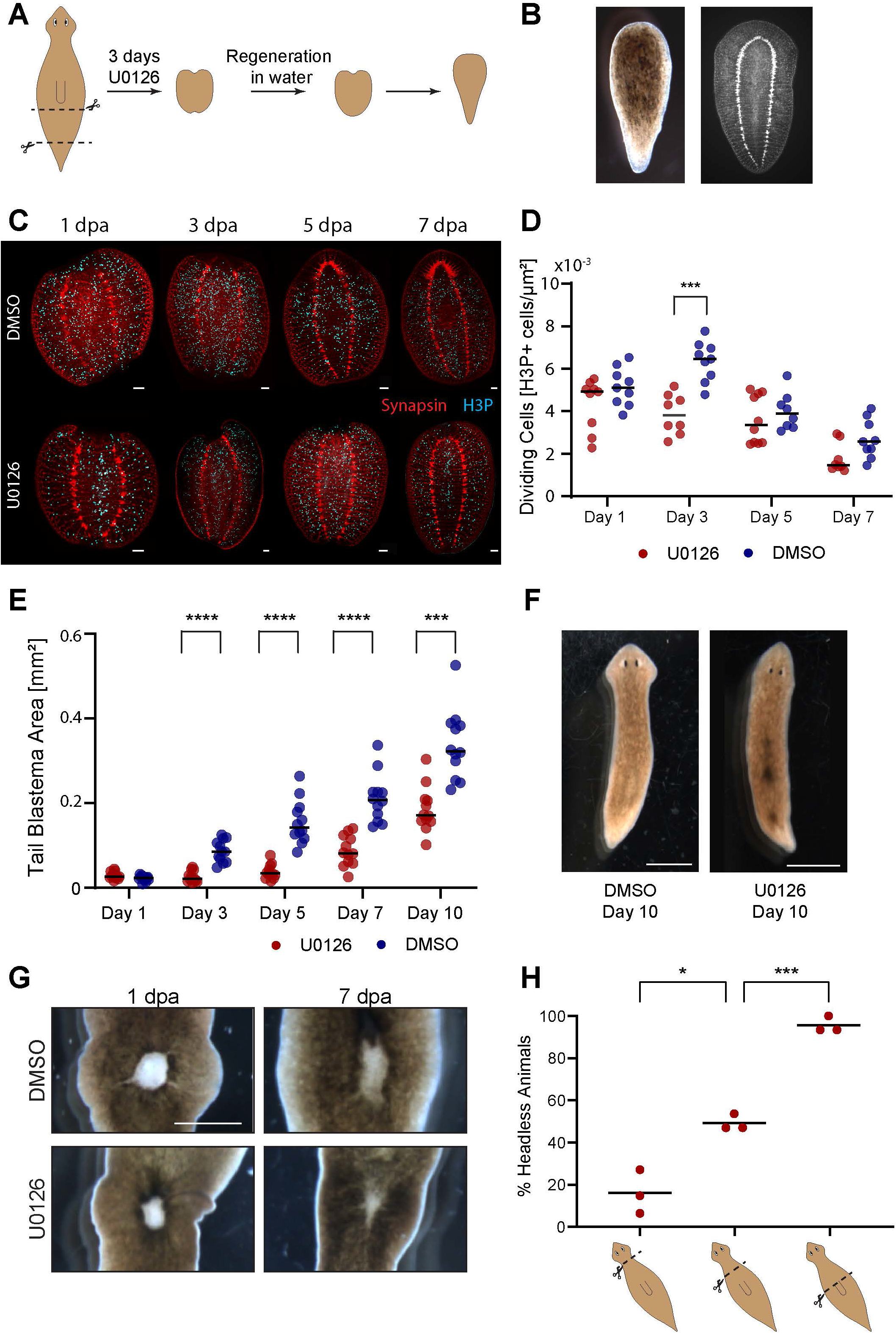
ERK inhibition blocks regeneration in a tissue- and location-specific manner. A) Experimental scheme showing the amputation planes of the pre-tail fragment treated with the ERK inhibitor U0126 for 3 days before completing regeneration in water. B) ERK inhibition in pre-tail fragments leads to the formation of headless animals, shown in both outside morphology and underlying neural structure via synapsin staining. C) Staining for synapsin (red) and dividing cells (H3P - cyan) in fragments treated with U0126 or DMSO as control, fixed at 1, 3, 5 and 7 dpa (days post amputation). Scale bar 100 μm. D) Quantification of the number of H3P foci/μm^2^ showing significant reduction of dividing cells in U0126 treated samples (red) compared to control samples (blue) at Day 3. N=10 E) Size of the tail blastema following amputation and treatment with U0126 or DMSO, showing significant difference at all timepoints after 3 dpa. N=12 F) Representative image of animals at 10 dpa after U0126 and control treatment, showing morphologically similar tail regeneration. Scale bar 500 μm. G) Progress of internal tissue regeneration following puncture wound at 1 dpa and 7 dpa in control and U0126 treated animals, showing similar progress of regeneration. Scale bar 500 μm. H) Percentage of headless animals formed following amputation at different planes along the anterior-posterior axis and U0126 treatment, showing significant differences between the amputation planes. Remaining worms regenerated as single-headed animals. N=3×15. *p< 0.05, *** p< 0.001, **** p<0.0001

The regeneration of the nervous system under ERK inhibition has not previously been described in detail. Therefore, we used staining for synapsin to show the formation of the neural anatomy of the headless planarians. This staining revealed that instead of the clusters of neural tissue that are visible at Day 3 in control animals’ anterior blastema, no staining indicative of brain formation was observed in U0126-treated samples. By Day 7 the two VNCs in the U0126-treated samples joined and form a rounded terminal structure (Figure 1C). These structures appeared to not form in the very small anterior blastema but rather in the underlying preexisting tissue.

ERK is known to play a crucial role in regulating cell division and we hypothesized that the lack of observed regeneration is due to a block of cell division. By staining for the cell division marker phosphohistone H3 (H3P), we observed that U0126-treated animals had significantly lower numbers of dividing cells at Day 3 compared to controls (Figure 1C, D), indicating a lack of the typical upregulation of cell division important in regeneration^18^. At the same time, there were still plentiful dividing cells even following 3 days of treatment with the highest tolerated dose of U0126.

The fact that dividing cells are observed even under the applied ERK inhibition may explain why, in addition to the clear lack of head regeneration, we found that the posterior wound of the fragments treated with U0126 formed a blastema and regenerated a normal tail shape (Figure 1B). To further investigate this apparent tissue-specific impact of ERK inhibition, we bisected animals and tracked tail regeneration on the anterior fragments following the 3-day treatment with U0126 (Figure 1E and F). The blastema area was significantly smaller in ERK inhibitor treated animals compared to DMSO treated controls from Day 3 to Day 10 (Figure 1E). But even though regeneration was reduced in U0126-treated animals, a clear increase in size of the tail blastema over time was apparent in the treated animals (Figure 1E) and the final regenerative outcomes at Day 10 were morphologically comparable to controls (Figure 1F). Similarly, animals treated with the ERK inhibitor for 3 days were able to replace sections of internal tissue lost through a puncture (Figure 1G). This indicates that while temporary ERK inhibition reduces cell division and delays regeneration, this regenerative delay can be overcome and normal regeneration can be resumed for the formation of all tissues except the head, which is unable to recover from the temporary block. This differential response in regeneration of different structures suggests a separate head-specific role of ERK signaling in addition to its established role in regulating cell division. This observation is consistent with the reports that while ERK is active in both wound sites immediately after amputation, it remains persistently active for 72h only in the anterior blastema^19^.

The ERK inhibition we used to induce the headless phenotype was achieved via a pharmacological agent, U0126, which offers the opportunity of washing out the inhibitor and therefore lift the inhibition. To test the time frame in which the inhibitor is removed from the tissue following our washout at 3 dpa, we first performed a mass spectroscopy analysis on planarian tissues incubated in U0126 for 3 days 1 day after washout and 7 days after washout. 1 day after the tissue was removed from the U0126 solution, only trace amounts of the inhibitor were detected (Supplemental Figure 1A). To confirm that those trace amounts were below the threshold at which they were affecting regeneration, worm fragments were placed in U0126 for 3 days, washed out and placed in regular worm water for 1 day before being re-amputated and allowed to regenerate (Supplemental Figure 1B). 92% of re-amputated animals regenerated as single-headed animals (n=45), indicating that the amount of U0126 remaining in the tissue at Day 3 was below the concentration necessary to inhibit head regeneration (Supplemental Figure 1B). These results indicate that following the washout after the 3 day incubation with U0126, the inhibition of ERK activity is lifted rapidly and suggests that the head-regeneration specific role of ERK signaling is required in the early regeneration time window of the first 72 hours post amputation, known to be critical for polarity setting^20,21^, and cannot be later replaced within the original regeneration time window without further injury to trigger the response.

### ERK inhibition blocks head formation in a location-specific manner

The different response of anterior and posterior blastemas to ERK inhibition led us to ask whether additional positional effects were present in the response to ERK inhibition. We therefore decapitated animal at different planes, ranging from directly behind the auricles in the head to above the pharynx, before treating the decapitated fragments with U0126 for 3 days and observing the regenerative outcomes (Figure 1H). Surprisingly, we found that decapitations just below the head results in normal regeneration following U0126 treatment in the vast majority of cases, while decapitation at the top of the pharynx resulted in almost entirely headless regeneration (Figure 1H). The intermediate cutting plane led to a mixed regenerative outcome with 51±4% single-headed animals while the remaining animals were headless (Figure 1H). Control animals regenerated as single-headed animals independent of cutting planes. This difference in regenerative outcomes is striking, as even the most anterior cutting plane requires reformation of the head with all its complex structures. The observed difference in regenerative ability along the A/P axis is reminiscent of the regeneration outcomes in animals treated with RNAi against the activin inhibitor Follistatin, which similarly showed headless regeneration only in more posterior cutting planes^11^. This parallel conforms with previous reports that suggests that ERK ERK activity is upstream of follistatin^7^ and indicates that ERK signaling may impact cell proliferation and differentiation differently in the context of a high Wnt signaling environment in the posterior compared to low Wnt anterior tissues. These connections between ERK and Wnt signaling are interesting in the light of studies showing interactions between these two signaling pathways in other systems^22^ and the hypothesis that ERK and Wnt/β-catenin form opposing gradients starting from head and tail respectively in planarians^9^.

### Some headless animals repattern to normal morphologies over long time

The headless animals that regenerate after ERK inhibition had previously been assumed to have reached a terminal, stable morphology^7^. We asked whether these animals would maintain their new morphology despite cell turnover or whether they would eventually be able to repattern to recover their original morphology (Figure 2A). Monitoring the headless animals for 18 weeks, we observed a remarkable phenotype in which some of the headless animals began to repattern, with some regaining a wild-type single-headed morphology. To quantify this phenomenon, we defined repatterning as any morphology change occurring after 4 weeks post-cutting. In headless animals observed from when they were cut until their death (n=181), 22% showed some form of repatterning (Figure 2B). Importantly, this repatterning occurred without any intervention. To ensure that no wounding happened to induce regeneration, animals were kept separated in individual wells and were not handled. No injury was observed during the weekly microscopic inspection of the animals. The remaining 78% of headless animals did not show any change in morphology except for shrinking in size, as headless animals are unable to feed and therefore have a limited lifetime. We observed repatterning from 4 weeks onwards with the majority of repatterning occurred between 4 and 10 weeks (Figure 2C). After 10 weeks only sporadic new repatterning was observed, although some worms began forming new heads as late as 18 weeks after cutting. This spontaneous repatterning that allows the re-establishment of a normal morphology many weeks after regeneration is completed suggests the existence of tissue processes that operate on a time scale much longer than previously known which trigger the re-emergence of normal body parts. The fact that we observe this repatterning, contrary to previous reports that suggested ERK-induced headlessness is a stable morphology, is most likely due to the longer timeframe of observation. Owlarn et al. state that headless animal remained headless even 18 days after amputation^7^, which does not contradicting our observation of repatterning 30+ days following amputation. Previous investigations of headless animals also mainly used the *Schmidtea mediterranea* species while we use *Dugesia japonica*, and the onset of repatterning may be distinct in the two species.

**Figure 2:**
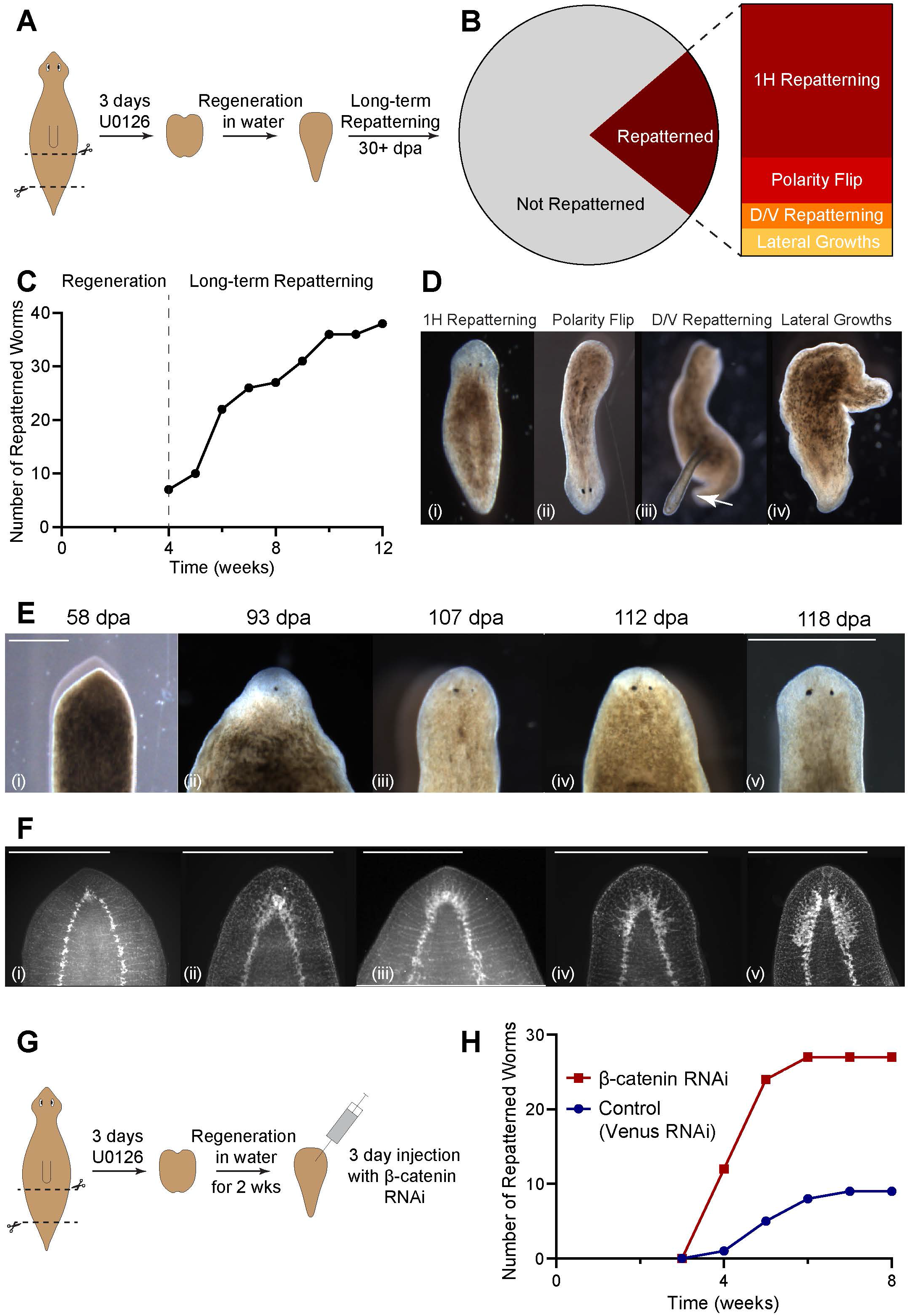
Headless animals can regain normal morphology over long timeframes. A) Illustration of the experimental procedure in which pre-tail fragments were treated for 3 days with U0126 to induce headless animals which were then observed for changes in morphology over long timeframes. B) Out of 181 headless worms tracked individual over time, 22% repatterned, while the remaining 78% did not change their morphology. The repatterned animals fell into 4 different categories – 61% Single-head (1H) repatterning, 18% polarity flip, 10% D/V repatterning, and 11% Lateral growths. C) Repatterning rate over time showing steady increase in the number of repatterned animals from 4 to 10 weeks after the initial amputation. D) Representative images of the 4 different repatterning types – i) 1H repatterning, ii) Polarity flip, iii) D/V repatterning (arrow marking dorsal outgrowth), and iv) Lateral growths. E) Repatterning to regain a single-headed morphology in one animal tracked over time. Scale bar 500 μm. F) Samples fixed and stained with synapsin antibody at different stages of repatterning, arranged into a putative neural repatterning timeline. Scale bar 500 μm. G) Experimental procedure with injection of β-catenin dsRNA or control dsRNA on 3 consecutive days into headless animals 14 days after initial induction. H) Repatterning rate in β-catenin RNAi (red) compared to Venus RNAi (blue) control animals. N=2×15.

Anatomically, the observed repatterning fell into 4 different categories. The most common repatterning type was single-headed repatterning where a normal wild-type morphology was regained (61% of all repatterning worms, Figure 2B, Di). Strikingly, the second most common phenotype was a polarity reversal, in which, following fissioning of a headless animal, the posterior blastema formed into a head (18%, Figure 2B, Dii). The remaining two categories represent the rare animals that exhibited growths which did not lead to the formation of a head, either in the form of dorsal outgrowths (D/V repatterning, 10%, Figure 2Diii) or lateral outgrowths (11%, Figure 2Div).

When headless animals repatterned to form a new head, the pigmentation of the round anterior end began to lighten and the tissue flattened out, forming a structure resembling a blastema (Figure 2E). Once this "blastema" formed, first one eyespot appeared (Figure 2E ii) and then the second one formed with a few days delay (Figure 2E iii), along with the head reshaping to regain the typical triangular morphology (Figure 2E iv and v). Animals were stained using synapsin antibodies to visualize the regrowth of the underlying brain tissue at intervals during the repatterning process. This showed that early brain repatterning occurred via the formation of a cluster of neural tissue at the anterior rounding of the VNC (Figure 2F i). Brain structures formed progressively from there, first in an apparently unorganized manner (Figure 2F ii), before expanding and reforming into a well-organized brain resembling that of a normal animal (Figure 2F iii-v). While the repatterning process resembles normal head regeneration in its progression, the timeframe was distinctly slower than normal head regeneration^23^, taking up to 25 days from the first observation of changes at the anterior to a fully formed head.

The observation that planarians are capable of recovering a wild-type bodyplan by spontaneously triggering a regeneration-like process, after maintaining a stable abnormal morphology for long periods of time, poses interesting questions about how abnormal morphologies are first maintained and how they are eventually detected to trigger correction. In other organisms there is some evidence that remodeling of tissues to fit new morphologies can be part of regeneration, such as the remodeling of transplanted tail tissue into a limb in amphibia^24^. The repatterning reported here is distinct from previous studies where it was shown that general injury signals can induce regeneration in headless animals^7^, as here there is no triggering injury of any kind. The here demonstrated ability to mount a regenerative response without any injury to trigger it is consistent with the idea that a missing tissue response (MTR) may not be necessary for regeneration to occur^11^. Our finding that the spontaneous repatterning is slower than normal regeneration is also consistent with other examples of regeneration in the absence of the MTR^11^.

The timepoint at which repatterning starts appears to be stochastic within the population. We showed that size was not a trigger for repatterning, as fragments of very different sizes could start repatterning at similar times (Supplemental Figure 2). This lack of correlation between repatterning size and timing rules out the starvation-induced shrinking as a trigger for repatterning.

Given that Wnt/β-catenin signaling is crucial in regulating head regeneration in planarians^25^, we investigated whether blocking Wnt signaling is able to drive repatterning in headless worms by injecting headless planaria with β-catenin dsRNAi and tracking their repatterning rates compared to headless animals injected with control dsRNAi (Figure 2G). Even though injection causes minor injury, we chose this method of dsRNAi administration due to the fact that headless planaria are unable to feed to ingest dsRNAi. We controlled for the injection-induced injury by the injection of a control dsRNAi, which led to a repatterning rate very similar to non-injected headless animals. In β-catenin RNAi animals, we saw repatterning of 100% of headless animals by 6 weeks after initial amputation, which was significantly more than in control RNAi-injected animals, where only around 30% repatterned in the same timeframe (Figure 2H). This suggests that repatterning uses the same signaling pathways that regulate normal head regeneration and that the start of repatterning may be dependent on Wnt signaling activity.

### Headless worms can reverse polarity following posterior injury

One of the unexpected repatterning phenotypes we observed was a polarity reversal, which occurred following spontaneous fissioning of headless animals. We observed fissioning even in headless animals that had not fed for a long time, showing that in *Dugesia japonica* fissioning can take place in the absence of a head and under low nutrition conditions. Following fissioning, the tail fragment always regenerated as a single-headed animal, as here the fissioning wound recapitulates an injury inducing head formation (Supplemental Figure 1). However, the anterior fragment has a more interesting, stochastic fate. In 85% of cases (82/96 observed fissioned animals), the anterior fissioning fragment regenerated the lost tail and remained headless. However, in the remaining 15% of cases (n=14/96) the anterior fissioning fragment underwent regeneration in which the posterior blastema formed into a head rather than a tail (Figure 3A). This resulted in the formation of single-headed animals with a head on the previously posterior end. In those animals, the rounded, formerly anterior, end repatterned into a typical pointed tail shape (Figure 3A iii-v). The single-headed animals arising from this polarity reversal exhibited striking movement patterns, in which the tail led the direction of the movement as opposed to the head in normal animals (Supplemental Movie 1). This abnormal movement confirms the reversal of polarity and is in line with the observed slow adaptation of cilia orientation in newly formed double-headed animals to their new morphology^26^. A spontaneous polarity reversal has not been reported in planaria, although previously worked induced head growth at the posterior blastema through application of electrical currents^27^.

**Figure 3:**
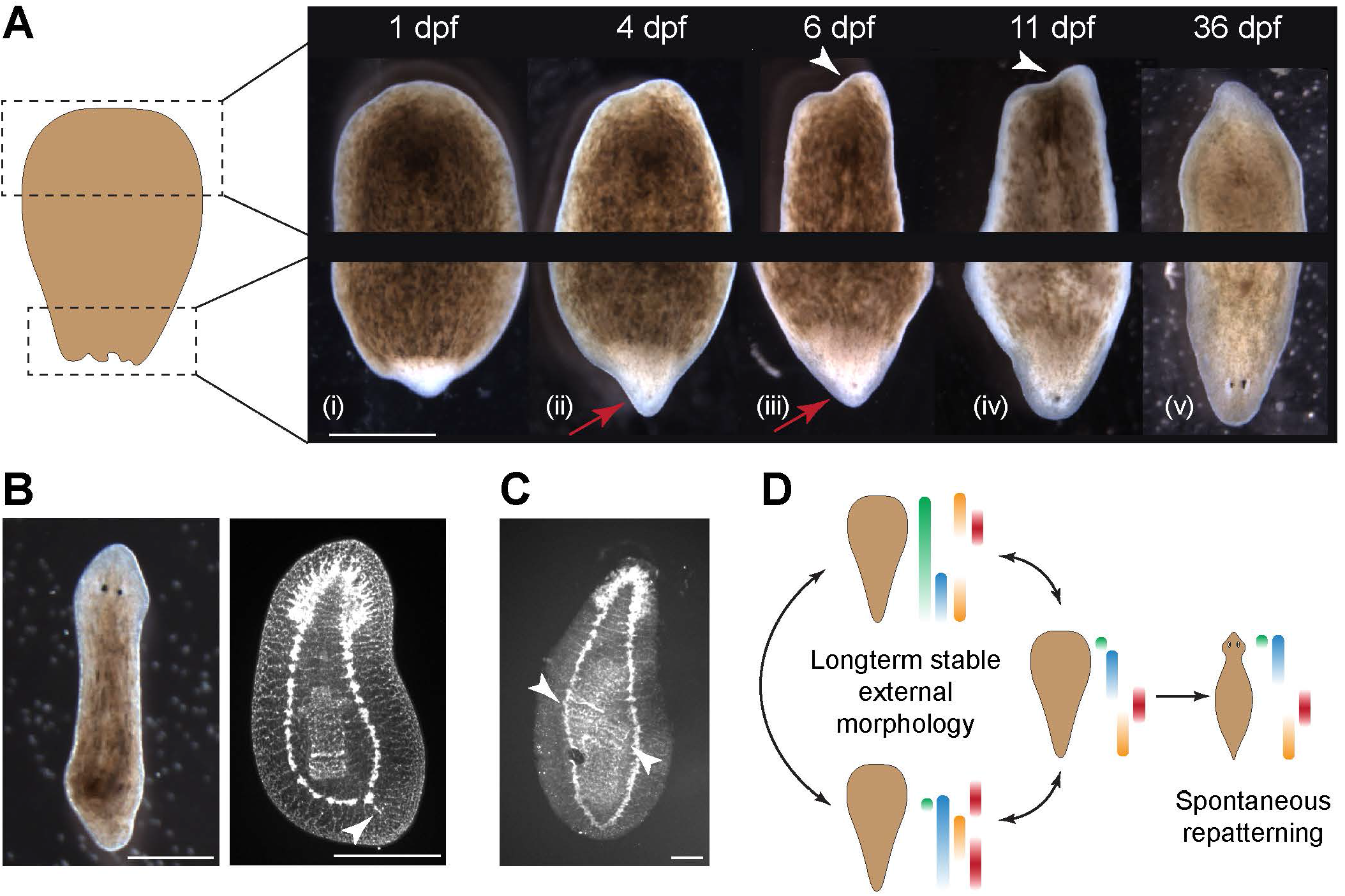
Headless worms can reverse their polarity. A) In the anterior fissioning fragment of a headless animal, the posterior blastema regenerates into a head, including the formation of an eyespot at 4 days after fissioning (red arrows). At the same time, the rounded anterior changes its morphology to a more typical tail-like morphology (white arrowheads). B) A single-headed worm resulting from polarity reversal in brightfield and synapsin staining of the same animal, illustrating normal brain morphology formed in the posterior blastema and beginning of changes in the VNC structure in the new tail (white arrowhead). Scale bar 500 μm C) Synapsin staining of a single-headed animal arising from polarity reversal with two pharynxes, the upper in the correct head-tail orientation and the lower in the opposite orientation, as visible by the distinct band and denser innervation around the pharynx mouth (white arrow). Scale bar 100 μm. D) Scheme illustrating the hypothesized misexpression of PCG domains that drives spontaneous repatterning of headless animals and polarity reversal.

This fascinating observation of a reversal of polarity following fissioning led us to investigate whether this reversal of polarity could also be induced by a posterior cut in headless animals. We therefore bisected headless animals at 4 weeks after induction at various planes along the A/P axis and observed the regenerative outcomes in the anterior fragments of all these cuts. Polarity reversal occurred in fragments from all different cutting planes, although at highly variable rates between experiments independent of the cut position (Supplemental Figure 3).

In animals that reversed their polarity, the nervous system regeneration went along with external morphology, with a normal brain forming in the posterior blastema (Figure 3B). At the same time, the formerly rounded VNC terminus began to branch out to reform the more pointed VNC tail morphology (Figure 3B). This however happened with a delay compared to the outward morphology, as worms with a relatively normal outward tail morphology were observed with the underlying nervous system only beginning to branch out (Figure 3B). Other internal structures also adjust to the new organismal polarity, often leading to abnormal intermediate states, such as worms with two pharynxes with opposing orientations (Figure 3C).

At the same time, many headless animals showed abnormal pharynx placement and orientation even before repatterning (Supplemental Figure 4). Most headless animals possessed pharynges at the very anterior end, directly abutting the rounded VNC anterior (Supplemental Figure 4A). At the same time, headless animals existed with multiple pharynges, either oriented perpendicular to each other and to the body axis (Supplemental Figure 3B) or aligned with the body axis but in opposing orientations to each other (Supplemental Figure 3C). The presence of animals with multiple and mispositioned pharynges may be indicative of a generally disturbed underlying whole-body axial polarity in headless animals, which results in a failure of signals about number and orientation of organ-level structures, leading to the disrupted formation of the pharynxes in location and orientation^28^.

Combining the observation of the polarity reversal at posterior injuries independent of A/P position with the identification of misplaced pharynges suggests a hypothesis explaining headless repatterning (Figure 3D). Headless animals may exhibit disorganized expression of the position control genes (PCGs) which control both regenerative outcomes and tissue maintenance^29^. Such a disorganized expression of regulating factors would explain why in a limited number of cases a posterior blastema gives rise to a head and why pharynges are formed in incorrect locations and positions. If the expression domains of the PCGs are not only mispositioned but also dynamically changing, this would explain the spontaneous starting times of repatterning of headless animals as resulting from the random alignment of a sufficient number of signaling factors to trigger tissue growth and formation of correct structures.

## Conclusion

ERK signaling is crucial in regulating the cell division required for regeneration. We show here that in planarian flatworms, ERK inhibition during the first 3 days of regeneration can be overcome in all tissues, except in regeneration of the head, suggesting a secondary signaling function for ERK in head formation. In the context of head regeneration, we present data that suggests that ERK signaling interacts with other signaling pathways, as ERK inhibition only blocks head regeneration in the posterior tissues which have high Wnt signaling activity, arguing for an interaction of ERK and Wnt signaling in regulating head regeneration.

We further report the surprising finding that the headless animals that form following ERK inhibition do not remain permanently headless, but over very long timeframes spontaneously begin to repattern and regain their normal morphology. In some of the headless animals, repatterning goes along with reversal of prior anterior-posterior axial polarity, suggesting an underlying instability of pattern signaling. The very long timeframe of the repatterning process (up to 18 weeks) poses interesting questions about what factors or physiological processes can be responsible for sensing mispatterning or triggering remodeling in this timeframe. It also suggests that other planarian regeneration outcomes should be observed over much longer times than is currently practiced, as other fascinating long-term effects may be discovered through this.

## Methods

### Colony maintenance

*Dugesia japonica* were maintained in Poland Spring water at 20°C, fed calf liver paste once a week and cleaned twice a week, as described in^30^. Animals were starved for one week prior to usage in amputation and pharmacological treatment experiments and were not fed for the duration of all experiments.

### Animal manipulation

Cutting of planaria was performed on a cooling plate using scalpel fragments. For generating headless animals, pre-tail fragments were cut by placing a cut at the pharynx opening and another cut narrowly above the tail tip. For tracking tail regeneration, animals were cut halfway between the head and tail. For tracking regeneration along the A/P axis, animals were either decapitated narrowly by cutting just below the auricles, cut halfway between the base of the head and the top of the pharynx, or cut directly at the top of the pharynx. To track regeneration of internal tissue a puncture wound was induced using square glass capillaries of a 0.7 mm^2^ inner diameter (VitroCom, Mountain Lakes, NJ) directly posterior to the pharynx opening. A fine paintbrush was used to remove the cut tissue from the middle of the animal.

### Pharmacological Treatments

Headless worms were produced through transient pharmacological treatment of freshly cut pre-tail fragments in 18 μM U0126 dissolved in DMSO (Sigma). No more than 40 fragments were treated per 10 cm petri dish. Fragments were incubated in U0126 for 3 days at 20°C before washing out the drug solution. At 14 days post amputation, the fragments were scored for regenerative phenotype.

### Scoring of Repatterned Phenotypes

Headless animals were maintained in individual wells of 12-well plates and their phenotypes were scored weekly for the duration of the experiment. Any significant changes in morphology, such as head regeneration, fissioning, or ectopic tissue development were noted. Time of repatterning was determines once one eye was visible in the forming head structure. Worms were labelled as “Polarity Flip” when a headless worm fissioned and a head regenerated at the posterior wound site. Worms were labelled as “Dorsal/Ventral repatterning” when an outgrowth of tissue occurred in the D/V plane of the headless worm. Worms were labelled as “Lateral growth” when significant changes in morphology occurred with growth on the lateral area of the animal, resulting in stable morphology which did not produce a head.

### RNA Interference

For β-catenin RNAi treatments, headless animals were selected at 14 dpa and injected on 3 consecutive days with dsRNA for *D. japonica* β-catenin and control dsRNA for VenusGFP. dsRNA was generated and injected as noted in^31^. 3 pulses of RNAi/per worm were given each day for 3 consecutive days.

### Mass Spectroscopy analysis of U0126 levels

Samples for MassSpec were generated by collecting fragments either directly from U0126 solution or following washout, removing all water, and placing 3 mm glass beads (Milipore, Burlington, MA) into an Eppendorf tube. Tissue was disrupted by vortexing for 2 min. DMSO was added as a solvent to the disrupted tissue and samples were vortexed again for 1 min. Liquid was removed from the beads and centrifuged for 20 min at 14,000 rotations per minute at 4°C. The upper clear phase was removed and filtered through a 0.2 μm polytetrafluoroethylene-syringe filter (Whatman, Maidstone, UK). Samples were stored at −80°C before analysis. U0126 standard was prepared by diluting in DMSO.

The samples were analyzed by the Harvard Faculty of Arts and Sciences Core Facility, on a Bruker Impact HD q-TOF mass spectrometer. Each LC-MS run was calibrated with known m/z values from sodium formate clusters. The LC column used was a henomenex Kinetex C18 (150 mm, 2.1 mm ID, 2.6 μm particle size). The HPLC method used 0.1% formic acid as mobile phase A and 0.1% formic acid in acetonitrile as mobile phase B. 5% B was used for the first 2 minutes, while the eluent was diverted to waste using a divert valve. At 2 minutes, the eluent was sent to the mass spectrometer. The mobile phase composition was changed linearly from 5% B to 100% B over 10 minutes. The composition was returned to 5% B over 0.1 minutes and kept constant for another 3.9 minutes to re-equilibrate the system to starting condition. A constant flow rate of 200 μL/min and an injection volume of 5 μL was used for all samples.

### Immunohistochemistry and Imaging

Worms were fixed in Carnoy’s fixative (2% HCl treatment to remove mucous, followed by fixing solution composed of 60% Ethanol, 30% Chloroform, and 10% Glacial Acetic Acid) and stained using primary antibodies anti-synapsin (SYNORF-1) (Developmental Studies Hybridoma Bank (DSHB), University of Iowa) at 1:50 and anti-phosphorylated histone H3 (H3P) (Invitrogen) at 1:250 dilution. Anti-SYNORF1 was deposited to the DSHB by Buchner, E. (DSHB Hybridoma Product 3C11 (anti SYNORF1))^32^. Secondary antibodies used were goat-anti-mouse Alexa488 (Sigma, St. Louis, MO, USA) and a goat-anti-rabbit-Alexa555 (Sigma) at 1:400 dilution. Samples were mounted in Vectashield^®^ hard set mounting medium (Vector Laboratories, Burlingame, CA) and imaged on a Nikon AZ100M stereoscope or a Leica SP8 confocal (Leica, Mannheim, Germany) with HyD detector, 488 and 552 nm diode light source, and 10x NA=0.4 (Leica HC PL APO CS2) objective.

### Image and Data analysis

All image analysis was performed in Fiji ImageJ^33,34^. To detect H3P positive cells, samples were thresholded using the Moments method, and the Analyze Particle function was used to count spot above a size of 10 pixels. All other analysis was performed manually. Data visualization and statistical analysis was performed in GraphPad Prism 8.2.1. ANOVA with Bonferroni post-correction were performed to determine significance.

### Data availability

All raw data is available upon request.

## Supplementary Materials

**Supplemental Figure 1:**
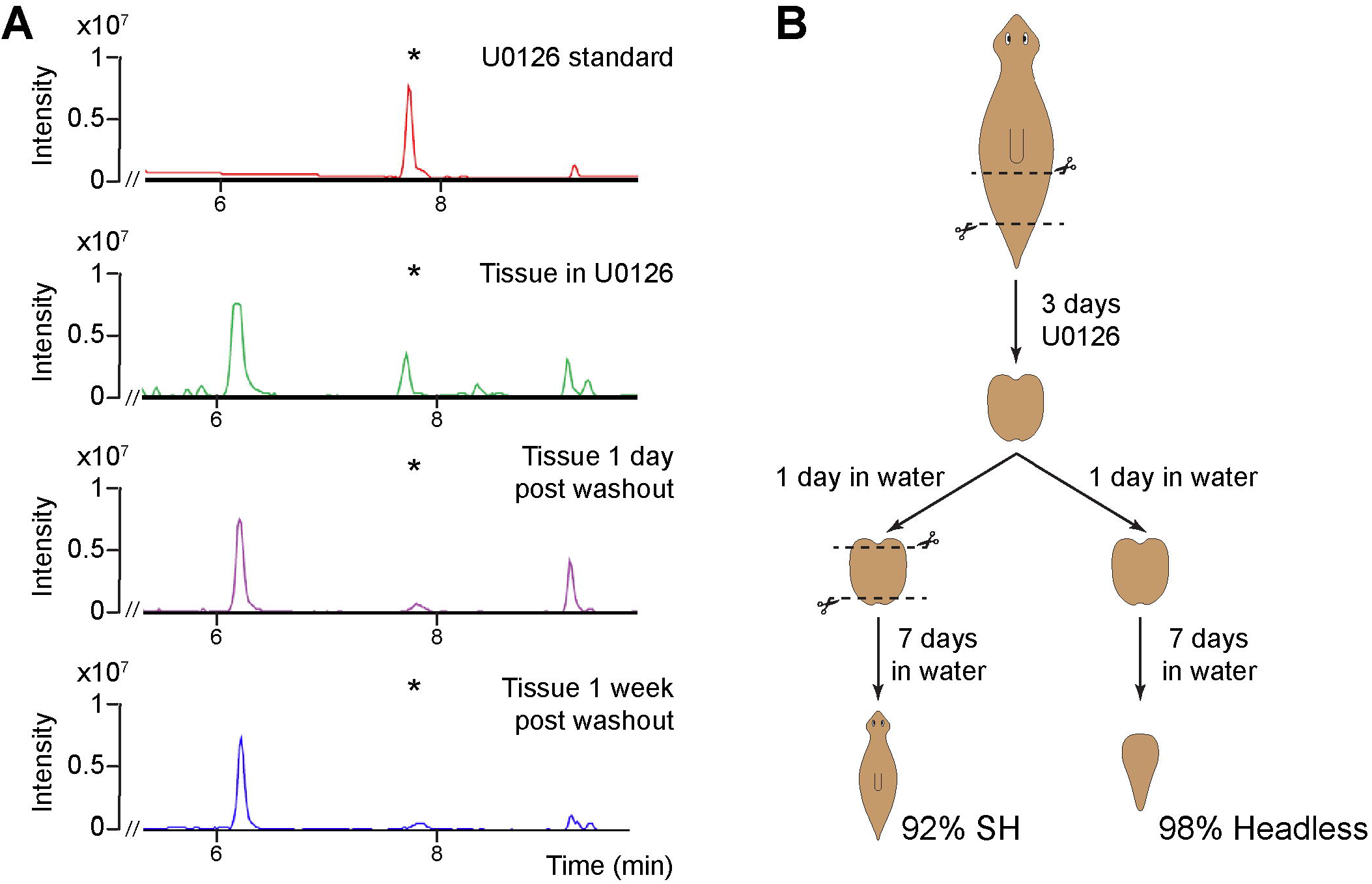
U0126 absence in tissue after washout. A) Mass spectroscopy analysis of a U0126 standard compared to extracts from planarian tissue incubated with U0126, tissue 1 day after washout from U0126 and 1 week after washout. Asterisk marks the characteristic U0126 peak. B) To show that the levels of U0126 remaining in the tissue 1 day after washout were not sufficient to inhibit regeneration, fragments were recut at 1 day after washout from U0126 before being allowed to regenerate. N=3 repeats of 15 animals each.

**Supplemental Figure 2:**
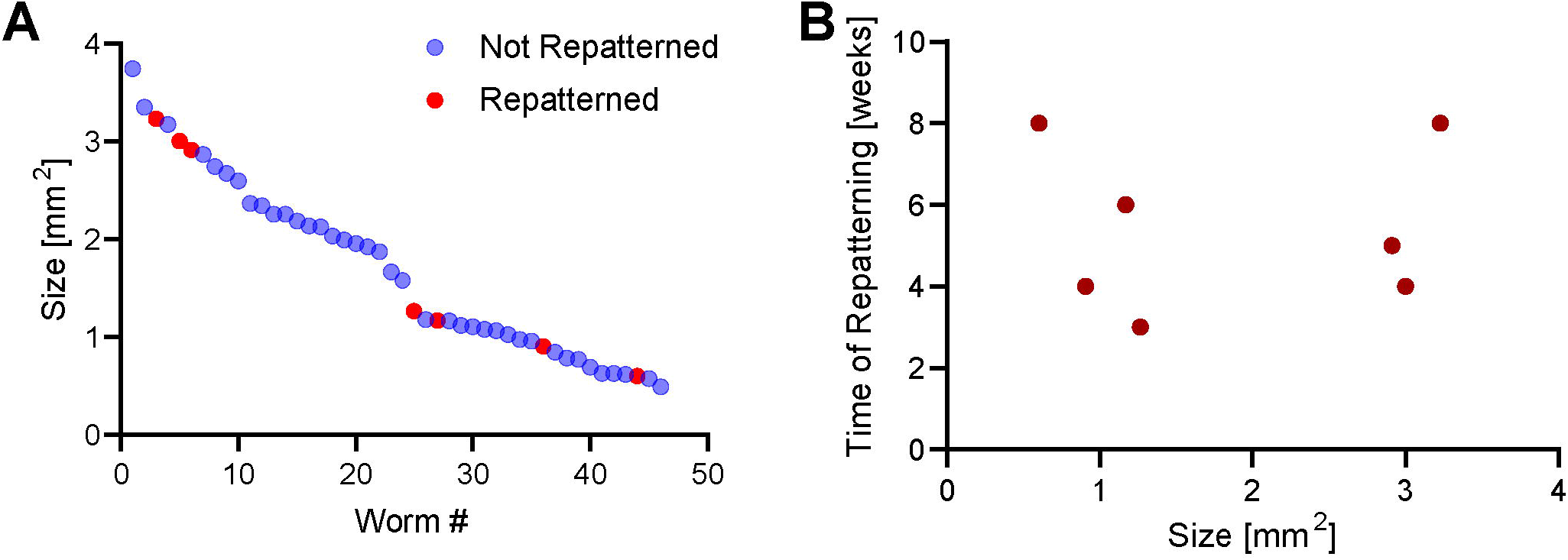
Timepoint of repatterning is independent of animal size. A) Size of all animals followed in this experiment showing wide range of sizes. Red points mark any animal that repatterned within the 8-week experimental timeframe. B) Repatterning timepoint plotted against size of the repatterned animal at the start of the experiment.

**Supplemental Figure 3:**
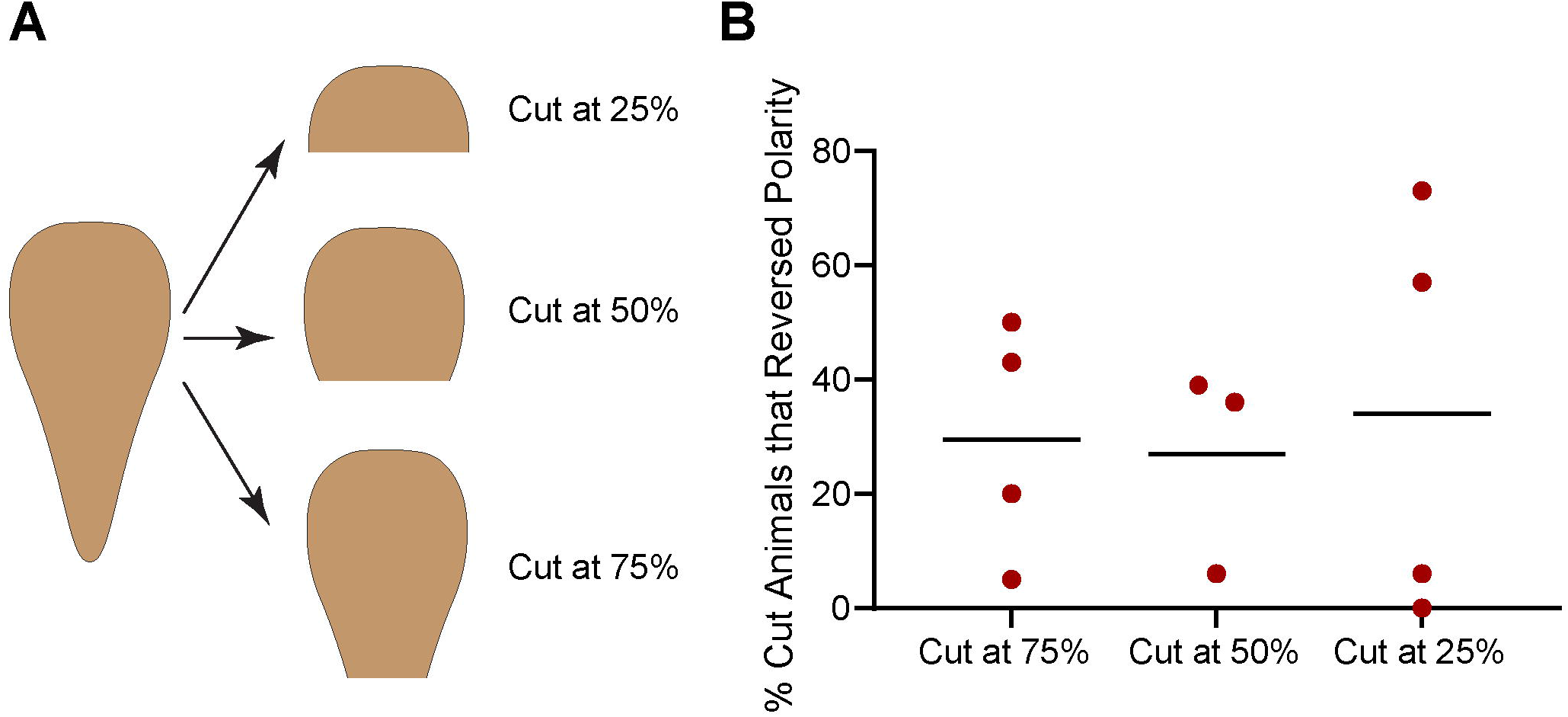
Polarity reversal results from cutting at variable rates. A) Experimental scheme showing the different planes at which the headless animals were cut. B) Percentage of animals that underwent polarity reversal following cuts at the respective planes. Each data point represents an independent repeat with 15 animals each.

**Supplemental Figure 4:**
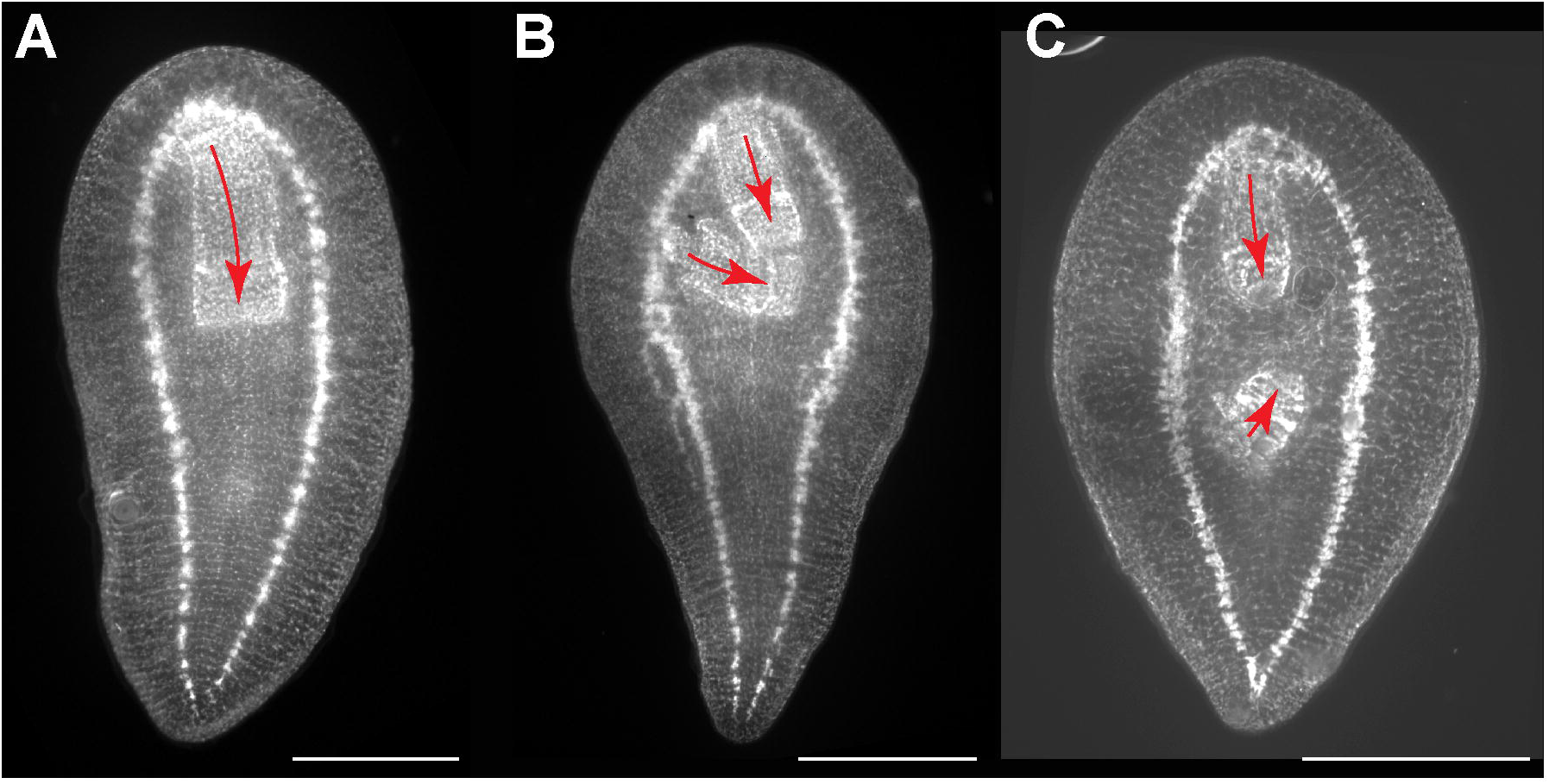
Synapsin stain of headless animals illustrating different pharynx positions. A) Headless animal with an anteriorly positioned pharynx. B) Headless animal with two pharynxes, one perpendicular to the head-tail axis. C) Headless animal with two pharynxes in opposing orientations. Scale bar 500 μm. Arrows illustrate pharynx orientation.

**Supplementary Video 1: Single-headed animal resulting from polarity reversal freely moving.**

## Acknowledgements

The authors would like to thank Anna Kane, Joshua Finkelstein, and all members of the Levin lab for thoughtful discussions on this project. We thank Hans Gonzembach for planaria colony maintenance. This research was supported by the Allen Discovery Center program through The Paul G. Allen Frontiers Group (12171), and by the National Institutes of Health Research Infrastructure grant NIH S10 OD021624. We also gratefully acknowledge support of the Barton Family Foundation and the Elisabeth Giaque Trust.

## Author contribution

Conceptualization: JB, JVL, CF, ML

Methodology Development: JB, JVL

Validation: JB, KAM

Formal Analysis: JB

Investigation: JB, JVL, KAM, KBW, JM

Resources Provision: ML

Data Curation: JB

Writing – Original Draft: JB

Writing – Review & Editing: JB, JVL, KAM, KBW, JM, CF, ML

Visualization: JB, JVL

Supervision: JB, ML

Project Administration: JB, ML

Funding Acquisition: ML

